# rplec: An R package of placental epigenetic clock to estimate aging by DNA-methylation-based gestational age

**DOI:** 10.1101/2025.02.04.636367

**Authors:** Herdiantri Sufriyana, Emily Chia-Yu Su

## Abstract

**Background:** Latest placental epigenetic clocks (PlECs) were claimed to be robust when applied to cases with either maternal or fetal adverse conditions. However, the accuracies in estimating gestational age (GA) were lower in earlier trimesters. We have developed a multistage predictive model to improve the accuracies, but it resulted in a large and complex PlEC, which may be less usable for non-computational scientists. To improve the usability of our PlEC, we aimed to develop an R package of PlEC to estimate aging by DNA-methylation-based GA (DNAm-GA).

**Methods:** An R package was developed to simplify our scikit-learn models into a single function and to utilize DNAm-GA for placental aging study. We provided two functions to normalize DNA methylation values and estimate DNAm-GA. Both were designed to run such that a user can adjust the number of samples per batch to fit their computational resources. Our model was simplified into a simple additive operation to reduce the need for expensive computation. Furthermore, two functions to perform quality control and identify placental aging. Quality control is performed by root mean squared-error (RMSE), mean absolute difference, and correlation coefficient. We defined placental aging as the deviation of placental DNAm-GA from the true GA beyond that from the measurement error.

**Results:** The simplified version of PlEC achieved similar performance with the original scikit-learn model with RMSE 0.102 (95% CI 0.101, 0.104), which was reasonably imperfect since Python and R handle floating/decimal numbers, differently. In our use case example, we could observe significant difference of placental aging in a specific period between case and control.

**Conclusions:** Our R package could reduce the computational requirement to use our models and maintained the precision in estimating DNAm-GA and our analytical framework could utilize DNAm-GA for placental aging study. Our PlEC also allows individual assessment of placental aging in clinical settings via the residual DNAm-GA.

## Introduction

Epigenetic clock is emerging in the last decade, as a tool for identifying whether a medical condition involves an accelerated aging [1-3]. It is also considered as having a pivotal role in placental pathology [4], leading to great obstetrical syndromes that result in preterm delivery [5]. While placenta is inaccessible before termination of pregnancy, placental epigenetic clock (PlEC) for clinical application is more feasible via maternal blood because of the advancement of cell-free fetal DNA [6, 7]. However, currently available PlECs are less precise for placentas from pregnant women who underwent preterm delivery.

Placental pathology plays a putative role in great obstetrical syndromes [8, 9]. They consist of preeclampsia, fetal growth restriction, preterm labor, preterm premature rupture of the membranes, late spontaneous abortion, and placental abruption [10]. These conditions lead to earlier pregnancy termination up to 10. 6% (95% confidence interval [CI] 9.0, 12.0) because of either spontaneous preterm delivery or medically-indicated termination [11]. The conditions contribute both maternal mortality and neonatal morbidity [12, 13]. The latter likely result in admission to neonatal intensive care unit. Although the conditions have a devastating impact in maternal and child health, a hallmark of placental pathology, that causes those conditions, is still unclear [14]. Placental aging role may be central in their pathophysiological derangements. Understanding placental aging is helpful for discovering the effective therapeutic/preventive strategy [15]. Therefore, a widely accessible tool is needed for studying placental aging, including PlEC which is a tool that precisely estimate gestational age (GA) based on placental DNA methylation.

Latest PlEC were claimed to be robust when applied to cases with either maternal or fetal adverse conditions because of three reasons [16]. First, they included samples with these conditions to develop the placental clocks. Second, ones for control samples and uncomplicated term accurately estimated GA in complicated pregnancies. Third, correlations between DNA methylation levels and GA remained significant after adjusting for preeclampsia (PE). However, simply including pathological samples would not remove bias, particularly due to confounders and colliders [17-20]. The accuracy was also arguable since it was lower in earlier trimesters [16], including another placental clock from Mayne, Leemaqz [21]. Eventually, the confounding adjustment was insufficient since PE is not the only condition that might affect both placental methylation and GA [22-24]. Previous PlECs [16, 21] also did not correct selection bias, which might reduce the accuracy in population with different distributions of adverse pregnancy conditions.

Recently, the Dialogue for Reverse Engineering Assessment and Methods (DREAM) held the Placental Clock DREAM Challenge, a crowd-sourced competition to develop and validate a more precise PlEC [25, 26]. This challenge provided an independent test set, which avoid the participants to evaluate their model until the final model submission, by containerizing their model via Docker. We participated in this challenge and our prediction model was selected as the top performer according to the test set [27]. This model was developed by multistage predictive modeling which resulted in a large and complex PlEC. It may be less usable for non-computational scientists in medicine and molecular biology to identify placental aging in the condition of interest. To improve the usability of our PlEC, we aimed to develop an R package of PlEC to estimate aging by DNA-methylation-based GA (DNAm-GA).

## Methods

The details of predictive modeling are described in the challenge webpage [28]. To facilitate user, we developed an R package to use our PlEC which is a multistage prediction model. It consists of *2n + 5* prediction models where *n* was the number of phenotypes that were correlated to GA. The input must be normalized among the specific probes for each sample and each model has the specific parameters for scaling the input. Furthermore, the ultimate goal of PlEC is to study placental aging in a condition of interest, which requires an understanding of how PlEC works and utilize its properties to identify accelerated or decelerated aging. Therefore, we need to achieve two objectives in developing this R package: (1) to simplify the usage of our model; and (2) to utilize DNAm-GA for placental aging studies.

### Simplifying the usage of our model

For the first objective, we need two functions to normalize DNA methylation values and estimate DNAm-GA. We need these functions to run per sample/thus, a user could run only a batch of samples each time. It is important to fit their computational resource (e.g., random-access memory [RAM] size).

The beta mixture quantile dilation (BMIQ) normalization is inherently performed for one sample across beta values at all the available CpG sites. In our model development, we applied BMIQ using *champ*.*norm* function from ChAMP R package. Its installation is quite challenging due to massive dependencies to other R packages. We created a function by modifying a part of *champ*.*norm*, that was specifically used for preparing input of our model. This function normalizes beta values per sample across beta values at the selected CpG sites specifically for our model.

The models in the multistage prediction model were originally developed using scikit-learn and other Python libraries. However, we need to simplify each model into a simple additive operation in R language to reduce the need for expensive computation and dependencies to the Python libraries. It is possible due to the nature of elastic net regression algorithm. The mathematical definition is shown for the multistage prediction model (Equation 1), where we developed the model to *α* estimate GA among normal samples, as denoted by 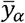. Deviation of 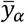 from the true GA, particularly among abnormal samples, is estimated by 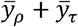.

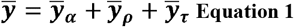

We developed the model *ρ* to estimate the first residual 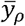 among samples across phenotypes *J* Where *j ∈ J* and each was *j* related to the true GA (Equation 2). The models *j*_1_ and *j*_2_ were respectively used for: (1) computing the probability of *j* occurring in a sample; and (2) estimating the residual among samples with phenotype *j*. Finally, we employed the model *ρ* to estimate 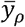. If the true GA is deviated from 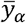, particularly among abnormal samples, then the residual is estimated by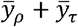. Any deviation of 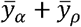 from the true GA may be contributed from other phenotypes beyond *J*.

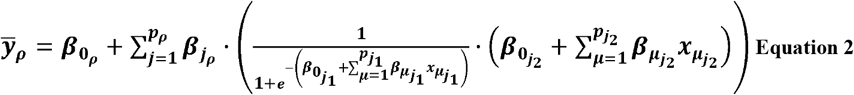

We developed the model *τ* to estimate the second residual 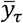 among samples across different GA periods of pregnancy termination (Equation 3). The placental samples were mostly, if not all, collected at the time of pregnancy termination which might be either spontaneous or medically-indicated. Based on domain knowledge, we considered that the pregnancy termination had different characteristics for preterm, term before the date, and term after the date. Respectively, the residual 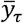 optimally estimated by 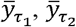, and 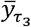. Earlier termination leading to preterm delivery is more likely due to phenotypes beyond *J*, that are correlated to the true GA.

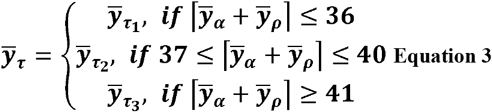

Either model *M* and *ρ* from scikit-learn in Python can be re-implemented using a simple mathematical operation in R (Equations 2 and 4). We only need to obtain each parameter *β* for predictor *i* from the scikit-learn model, and the average and standard deviation to scale the normalized beta value *x*_*i*_. Eventually, for technical validation, we compared 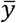 from the multistage prediction model in scikit-learn with 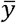 from the function in our R package.

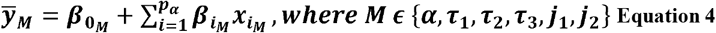

### Utilizing DNAm-GA for placental aging studies

For the second objective, we need two functions to perform quality control and identify placental aging. We need the former function to ensure our PlEC is well-generalized using the user’s own dataset. Meanwhile, the latter function is used to compare aging across GA between case and control groups in the same dataset.

Quality control is performed by several metrics. We provide a function to create calibration plot and compute root mean squared-error (RMSE), mean absolute difference (MAE), and Pearson’s correlation coefficient (*r*). A user is suggested for proceeding to placental aging identification if these metrics are similar to those in the validation set.

For identifying placental aging, we defined it as the deviation of placental DNAm-GA from beyond that from the true GA *y* beyond that from the measurement error *e* (Equation 5). If the error value is acceptable, then “abnormal” placental aging is the residual 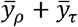, as denoted by 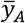. It is because we developed the model *α* to estimate placental aging 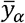 among normal samples. However, normal placenta may have accelerated/decelerated placental aging to some extent but not leading to abnormal condition.

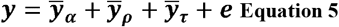

To determine whether placental aging is abnormal, the residual 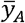 is compared between case and control groups. Specifically, the null hypothesis is defined in Equation 6 where 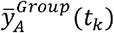 represents placental aging for each group at timepoint *t* as a percentile (*k* = [1,2,…, 100]) between *a* and *b* which are respectively the minimum and maximum GA across all groups in the user’s dataset. Accordingly, the alternative hypothesis is stated in Equation 7. If *H*_0_ is rejected, it indicates that placental aging is significantly different between case and control groups, suggesting abnormal placental aging in case group if the control represents the normal one.

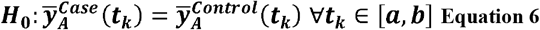

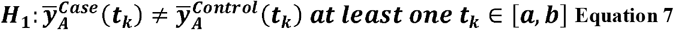

Notably, placental samples case and control groups are sampled at different GA. Meanwhile, the comparison is paired between case and control for every *t*_*k*_. Hence, we need to interpolate and extrapolate the residual 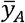 as a function *g*_*A*_ over a common range of *t* from *a* to *b* (Equation 8).

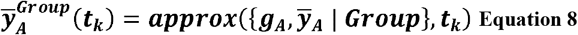

We also provide an aging plot which is a line plot where x-axis is GA in weeks and y-axis is case-control aging difference in weeks. A ribbon plot is shown to depict the uncertainty interval via bootstrapping. Furthermore, we allow a user to determine the values of *a* and *b* for testing the hypothesis within a specific range of GA that demonstrated accelerated/decelerated placental aging in the plot.

For use case examples, we randomly selected a sample from each combination of GA periods and preeclampsia status in the training set [28]. The GA period consisted of ≤12, 13 to 26, 27 to 36, 37 to 40, and >40 weeks’ gestation. This selection was intended to cover all trimesters before and after the ideal date of delivery (40 weeks’ gestation).

## Statistical analysis

A technical validation was conducted by comparing all the metrics of between original and simplified versions respectively using the scikit-learn model and *rplec* function. Meanwhile, in placental aging analysis, we provided either Mann-Whitney U or permutationtest for testing the hypothesis. To provide the uncertainty interval in the plot, we resample 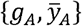 within each group for each bootstrap iteration to compute the standard deviation (SD). The ribbon ranges from 1 to -1 SD from 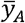. Therefore, a user can identify whether placental aging in case group is accelerated (>0) or decelerated (<0) compared to control group.

## Results

### Predictive performance of the simplified PlEC in *rplec*

Using the training set, we conducted technical validation for the simplified version of PlEC in our *rplec* R package. This version demonstrated similar predictive performance in all of the evaluation metrics with the original one (Figures 1A and 1B). The simplified version of PlEC achieved similar performance with the original scikit-learn model with RMSE 0.102 (95% CI 0.101, 0.104). A slightly different results between both version (Figure 1C) is reasonable since Python and R handle floating/decimal numbers, differently.

**Figure 1.**
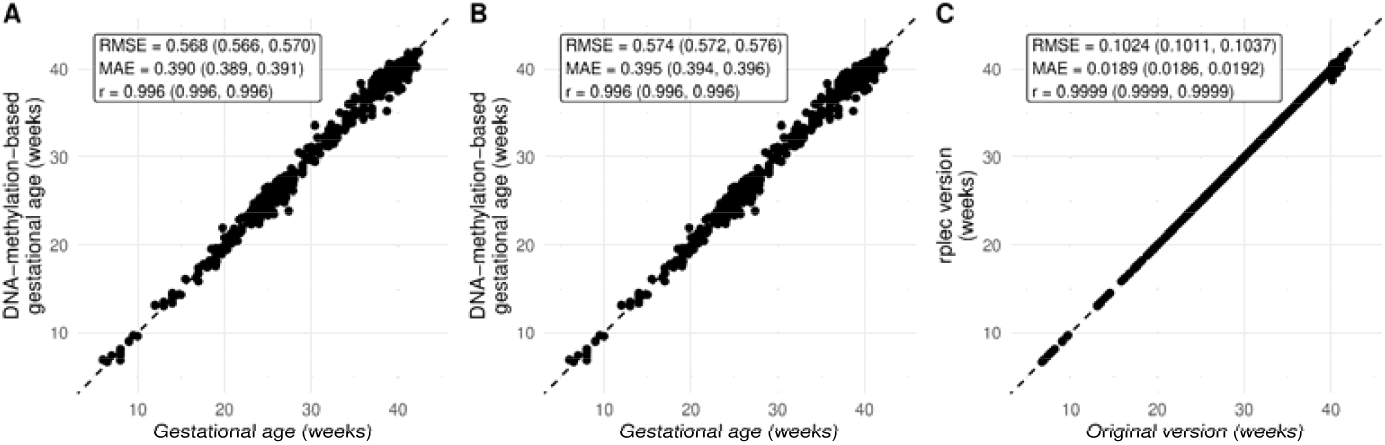
Technical validation of *rplec* function for GA estimation: (1) the original version using the scikit-learn model vs. the true GA; (2) the simplified version using *rplec* function vs. the true GA; and (3) the simplified version vs. the original version. MAE, mean absolute error (weeks); *r*, Pearson’s correlation coefficient; RPC, robust placental clock; RMSE, root mean squared-error (weeks).

### Use case example in utilizing DNAm-GA for placental aging studies

A user needs to install and load our R package either from the comprehensive R archive network (CRAN) or our GitHub repository (lines 1 to 3 Code snippet 1). We provided built-in data of beta values, GA, and phenotype (case/control) for 10 samples (lines 4 to 7 Code snippet 1), that were eligible as a use case (see “Utilizing DNAm-GA for placental aging studies”). After normalization and DNAm-GA estimation (lines 8 to 12 Code snippet 1), quality control is suggested to evaluate the predictive performance of our PlEC in the user’s own dataset (line 13 Code snippet 1). If the predictive performance is accepted (Figure 2A), then placental aging is estimated using the residual DNAm-GA (lines 14 to 15 Code snippet 1). Eventually, a user need examine whether placental aging is significantly different between case and control groups (lines 17 Code snippet 1).

**Figure 2.**
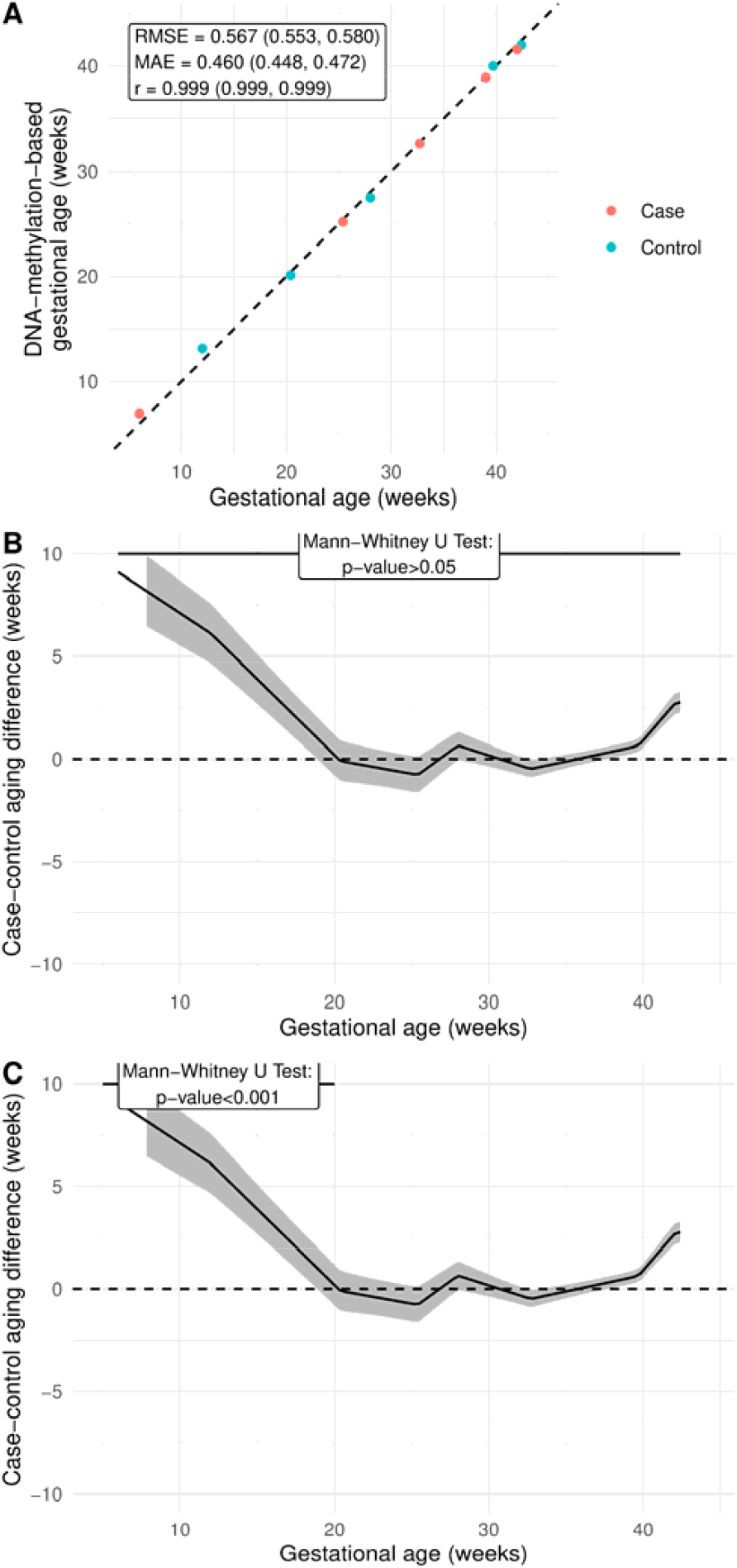
Placental aging analysis output in *rplec*: (A) quality control; (B) case-control aging difference with hypothesis testing across GA; and (C) case-control aging difference with hypothesis testing from 5 to 20 weeks’ gestation. MAE, mean absolute error (weeks); *r*, Pearson’s correlation coefficient; RMSE, root mean squared-error (weeks).

### Code snippet 1. Placental aging analysis

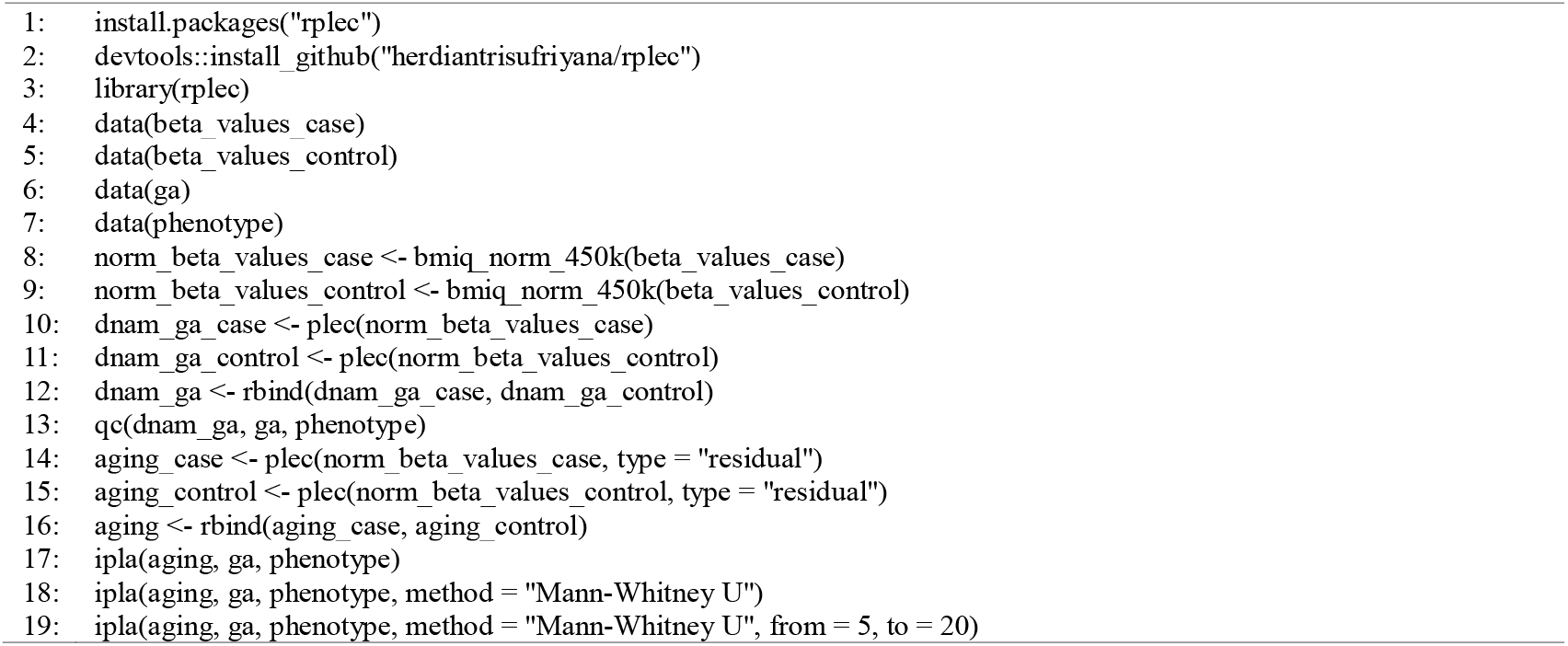

In this example (Table S1), we can observe difference in placental aging between case and control (Figures 2B and 2C). Using Mann-Whitney U test (line 18 Code snippet 1), placental aging in case group is not significantly different to that in control group across all trimesters (Figure 2B).

However, a larger difference is observed from ∼5 to ∼20 weeks’ gestation. In this period, placental aging in case group is significantly different to that in control group (Figure 2C; line 19 Code snippet 1). The y-axis indicates that the case group demonstrated an accelerated aging from ∼6 to ∼9 weeks within ∼8 to ∼12 weeks’ gestation. In other words, the placental age in this group was similar to ∼17 to ∼18 weeks’ gestation in control group during first trimester, but the accelerated aging is resolved after 20 weeks’ gestation.

## Discussion

### Finding summary

We have simplified our PlEC from 31 scikit-learn models into a single function in an R package, i.e., *rplec*, with a built-in function for preprocessing the input. The simplified version maintained the precision of the original model. We also provide another function to utilize our PlEC for a placental aging study, appropriately based on the design of our PlEC.

### Research and clinical implications

Generally, there are two main challenges in epigenetic clock research [3]. First, the challenge remains in identifying whether the acceleration is the precision error of a DNAm biological clock or a genuine biological aging effect. Second, biological age acceleration can be defined as: (1) absolute age acceleration (AAA), which is the difference between DNAm and chronological age; or (2) relative age acceleration (RAA), which is the residual of linear regression of DNAm age versus chronological age. In general, the second definition is preferred because it is arguably able to adjust sampling error. However, using RAA, the effect of age acceleration may not be detected on the outcome when it is used only in the case group, e.g., the effect on body mass index among individuals with clinical obesity [29].

Our modeling technique entangled the biological aging effect from the precision error (Equation 5). A user can re-evaluate the precision of our PlEC using their own dataset. The error may be technical or biological variation including that in the control group. Our statistical framework also corrects sampling error. It is because this framework only compares the residual DNAm-GA which refers to placental aging beyond those in normal aging process (Equations 6 and 7). Therefore, we can expect our PlEC and the statistical framework to refine current method for placental aging studies.

Several studies had used PlEC for placenta aging-related studies [30-32]. However, its clinical application is hampered regardless the advancement of cell-free fetal DNA, because RAA cannot be used to identify placental aging in a single sample. Our modeling technique allows the usage of the residual DNAm-GA for a single sample and comparison to those from the control samples at the same time. Hence, the residual DNAm-GA is expected to move forward the usage of PlEC for clinical application.

### Strength and limitation

Our study demonstrated that PlEC precision could be improved solely by modeling techniques beyond adding more data and applying more sophisticated algorithms. The residual DNAm-GA also offers individual assessment of placental aging based on AAA while also adjusts sampling error similar to RAA. Our PlEC can be re-evaluated in every placental aging study using quality control function in the R package.

However, we also considered several limitations in this study. First, a model in our PlEC may have variations of accuracy and precision respectively in classifying and estimating GA for a particular phenotype. This limitation may lead to poor precision when a user conducts quality control using their own dataset. Yet, the PlEC quality can be improved by partially retraining the model with more data for that phenotype. Second, our PlEC also did not include several phenotypes which might affect GA via early termination, e.g., placental abruption. Nevertheless, our modeling framework allows future studies to add a model and re-train some models in our PlEC. Third, our PlEC still required the whole array. While it can be a burden for clinical application, our PlEC is still useful for research purpose and reducing the predictors further with similar accuracy.

## Conclusions

We have developed an R package of PlEC to allow users to preprocess their data and conduct quality control using their own data. Eventually, we provide a function for a user utilize DNAm-GA for placental aging study. In addition, our PlEC allows individual assessment of placental aging in clinical settings.

## Acknowledgments

Preprint of an article published in the 47^th^ Annual International Conference of the IEEE Engineering in Medicine and Biology Society (EMBC). This study was funded by: (1) the Postdoctoral Accompanies Research Project from the National Science and Technology Council (NSTC) of Taiwan (grant nos.: NSTC111-2811-E-038-003-MY2 and NSTC113-2811-E-A49A-003) to HS; and (2) the National Science and Technology Council in Taiwan (grant no. NSTC113-2221-E-A49-193-MY3), the Ministry of Science and Technology (MOST) of Taiwan (grant nos.: MOST110-2628-E-038-001 and MOST111-2628-E-038-001-MY2), the University System of Taipei Joint Research Program (grant no.: USTP-NTOU-TMU-112-04), and the Higher Education Sprout Project from the Ministry of Education (MOE) of Taiwan (grant no.: DP2-111-21121-01-A-05 and DP2-TMU-112-A-13) to ECYS. These funding bodies had no role in the study design; in the collection, analysis, and interpretation of data; in the writing of the report; or in the decision to submit the article for publication.

## CRediT authorship

**HS:** Conceptualization, Methodology, Software, Validation, Formal analysis, Writing – original draft, Visualization, Funding acquisition. **ECYS:** Conceptualization, Methodology, Resources, Writing – review & editing, Supervision, Funding acquisition. All authors have read and approved the manuscript and agreed to be accountable for all aspects of the work in ensuring that questions related to the accuracy or integrity of any part of the work are appropriately investigated and resolved.

## Data availability

The R package can be installed either from the CRAN or our GitHub repository (https://github.com/herdiantrisufriyana/rplec). Data are publicly available in GEO where the accession numbers are provided in the source codes. However, the curated version (https://www.synapse.org/Synapse:syn59520082/wiki/628063) can only be accessed by request to the challenge organizer.

## References

1. Bocklandt, S., et al., Epigenetic predictor of age. PLoS One, 2011. 6(6): p. e14821.

2. Horvath, S. and K. Raj, DNA methylation-based biomarkers and the epigenetic clock theory of ageing. Nat Rev Genet, 2018. 19(6): p. 371–384.

3. Teschendorff, A.E. and S. Horvath, Epigenetic ageing clocks: statistical methods and emerging computational challenges. Nat Rev Genet, 2025.

4. Pan, M., et al., The role of placental aging in adverse pregnancy outcomes: A mitochondrial perspective. Life Sci, 2023. 329: p. 121924.

5. Ciampa, E.J., et al., Hypoxia-inducible factor 1 signaling drives placental aging and can provoke preterm labor. Elife, 2023. 12.

6. Goldwaser, T. and S. Klugman, Cell-free DNA for the detection of fetal aneuploidy. Fertil Steril, 2018. 109(2): p. 195–200.

7. Lo, Y.M.D., et al., Epigenetics, fragmentomics, and topology of cell-free DNA in liquid biopsies. Science, 2021. 372(6538).

8. Brosens, I., P. Puttemans, and G. Benagiano, Placental bed research: I. The placental bed: from spiral arteries remodeling to the great obstetrical syndromes. Am J Obstet Gynecol, 2019. 221(5): p. 437–456.

9. Harris, L.K., et al., Placental bed research: II. Functional and immunological investigations of the placental bed. Am J Obstet Gynecol, 2019. 221(5): p. 457–469.

10. Brosens, I., et al., The “Great Obstetrical Syndromes” are associated with disorders of deep placentation. Am J Obstet Gynecol, 2011. 204(3): p. 193–201.

11. Chawanpaiboon, S., et al., Global, regional, and national estimates of levels of preterm birth in 2014: a systematic review and modelling analysis. Lancet Glob Health, 2019. 7(1): p. e37–e46.

12. Say, L., et al., Global causes of maternal death: a WHO systematic analysis. Lancet Glob Health, 2014. 2(6): p. e323–33.

13. Saleem, S., et al., Neonatal deaths in infants born weighing[≥[2500 g in low and middle-income countries. Reprod Health, 2020. 17(Suppl 2): p. 158.

14. Hoffman, M.K., The great obstetrical syndromes and the placenta. Bjog, 2023. 130 Suppl 3(Suppl 3): p. 8–15.

15. Li, Y., et al., Molecular mechanisms of aging and anti-aging strategies. Cell Commun Signal, 2024. 22(1): p. 285.

16. Lee, Y., et al., Placental epigenetic clocks: estimating gestational age using placental DNA methylation levels. Aging (Albany NY), 2019. 11(12): p. 4238–4253.

17. Greifer, N. and E.A. Stuart, Matching Methods for Confounder Adjustment: An Addition to the Epidemiologist’s Toolbox. Epidemiol Rev, 2022. 43(1): p. 118–129.

18. Liu, T., et al., Can statistic adjustment of OR minimize the potential confounding bias for meta-analysis of case-control study? A secondary data analysis. BMC Med Res Methodol, 2017. 17(1): p. 179.

19. Holmberg, M.J. and L.W. Andersen, Collider Bias. Jama, 2022. 327(13): p. 1282–1283.

20. Digitale, J.C., et al., Key concepts in clinical epidemiology: collider-conditioning bias. J Clin Epidemiol, 2023. 161: p. 152–156.

21. Mayne, B.T., et al., Accelerated placental aging in early onset preeclampsia pregnancies identified by DNA methylation. Epigenomics, 2017. 9(3): p. 279–289.

22. Jiang, M., et al., A case control study of risk factors and neonatal outcomes of preterm birth. Taiwan J Obstet Gynecol, 2018. 57(6): p. 814–818.

23. Shi, D., et al., Placental DNA methylation analysis of selective fetal growth restriction in monochorionic twins reveals aberrant methylated CYP11A1 gene for fetal growth restriction. Faseb j, 2023. 37(10): p. e23207.

24. Konwar, C., et al., DNA methylation profiling of acute chorioamnionitis-associated placentas and fetal membranes: insights into epigenetic variation in spontaneous preterm births. Epigenetics Chromatin, 2018. 11(1): p. 63.

25. Stolovitzky, G., D. Monroe, and A. Califano, Dialogue on reverse-engineering assessment and methods: the DREAM of high-throughput pathway inference. Ann N Y Acad Sci, 2007. 1115: p. 1–22.

26. Meyer, P. and J. Saez-Rodriguez, Advances in systems biology modeling: 10 years of crowdsourcing DREAM challenges. Cell Syst, 2021. 12(6): p. 636–653.

27. Placental Clock DREAM Challenge. 2024 June 3rd, 2024]; Available from: https://www.synapse.org/Synapse:syn59520082/wiki/628063.

28. Sufriyana, H. and E.C.-Y. Su. Refining placental clock measured by the Infinium HumanMethylation-450/850 BeadChip arrays by correcting collider-restriction bias. 2024 March 8th, 2024]; Available from: https://www.synapse.org/Synapse:syn62407322/wiki/629409.

29. Horvath, S., et al., Obesity accelerates epigenetic aging of human liver. Proc Natl Acad Sci U S A, 2014. 111(43): p. 15538–43.

30. Polinski, K.J., et al., Epigenetic gestational age and the relationship with developmental milestones in early childhood. Hum Mol Genet, 2023. 32(9): p. 1565–1574.

31. Saeed, H., et al., Placental accelerated aging in antenatal depression. Am J Obstet Gynecol MFM, 2024. 6(1): p. 101237.

32. Tekola-Ayele, F., et al., Sex differences in the associations of placental epigenetic aging with fetal growth. Aging (Albany NY), 2019. 11(15): p. 5412–5432.

